# A Joint Deep Learning Model for Simultaneous Batch Effect Correction, Denoising and Clustering in Single-Cell Transcriptomics

**DOI:** 10.1101/2020.09.23.310003

**Authors:** Justin Lakkis, David Wang, Yuanchao Zhang, Gang Hu, Kui Wang, Huize Pan, Lyle Ungar, Muredach P. Reilly, Xiangjie Li, Mingyao Li

**Author notes:** Correspondence: Justin Lakkis, Xiangjie Li, Mingyao Li, Ph.D.

## Abstract

Recent development of single-cell RNA-seq (scRNA-seq) technologies has led to enormous biological discoveries. As the scale of scRNA-seq studies increases, a major challenge in analysis is batch effect, which is inevitable in studies involving human tissues. Most existing methods remove batch effect in a low-dimensional embedding space. Although useful for clustering, batch effect is still present in the gene expression space, leaving downstream gene-level analysis susceptible to batch effect. Recent studies have shown that batch effect correction in the gene expression space is much harder than in the embedding space. Popular methods such as Seurat3.0 rely on the mutual nearest neighbor (MNN) approach to remove batch effect in the gene expression space, but MNN can only analyze two batches at a time and it becomes computationally infeasible when the number of batches is large. Here we present CarDEC, a joint deep learning model that simultaneously clusters and denoises scRNA-seq data, while correcting batch effect both in the embedding and the gene expression space. Comprehensive evaluations spanning different species and tissues showed that CarDEC consistently outperforms scVI, DCA, and MNN. With CarDEC denoising, those non-highly variable genes offer as much signal for clustering as the highly variable genes, suggesting that CarDEC substantially boosted information content in scRNA-seq. We also showed that trajectory analysis using CarDEC’s denoised and batch corrected expression as input revealed marker genes and transcription factors that are otherwise obscured in the presence of batch effect. CarDEC is computationally fast, making it a desirable tool for large-scale scRNA-seq studies.

## Introduction

Single-cell RNA sequencing (scRNA-seq) analysis has substantially advanced our understanding of cellular heterogeneity and transformed biomedical research. However, the analysis of scRNA-seq data remains confounded by batch effects, which are inevitable in analyses of human tissue and are prevalent in many scRNA-seq studies in general^1,2^. Several methods have been developed to remove batch effect in scRNA-seq data analysis^3-10^. These methods can be divided into two categories: 1) batch correction in the low-dimensional embedding space, and 2) batch correction in the original gene expression space. Most published papers belong to the first category^3,7-10^. Although useful for profiling the overall characteristics of cells such as clustering and trajectory reconstruction, these methods cannot be used for downstream gene-level analysis like differential expression and co-expression analysis.

A recent benchmarking study has shown that correcting batch effect in the gene expression space is much more challenging than in the embedding space^11^. Popular methods such as Seurat 3.0^6^ rely on the mutual nearest neighbor (MNN) approach^5^ to remove batch effect in the gene expression space, but MNN can only analyze two batches at a time. Its performance is affected by the ordering in which batches are corrected and it quickly becomes computationally infeasible when the number of batches gets large. Moreover, our evaluations indicate that MNN performs poorly for removing batch effect for genes that are not highly variable. Another popular method, scVI, suffers from a similar issue in which the denoised gene expression is still susceptible to batch effect, particularly for those genes that are not highly variable. Non-highly variable genes represent the majority of genes in the genome, where batch effects constitute a larger fraction of variance in the transcriptome and are much harder to correct.

To address this gap in the literature, we present CarDEC (**<u>C</u>**ount **<u>a</u>**dapted **<u>r</u>**egularized **<u>D</u>**eep **<u>E</u>**mbedded **<u>C</u>**lustering), a joint deep learning framework for simultaneous batch effect correction, denoising, and clustering of scRNA-seq data. Rather than explicitly modeling batch effect, CarDEC jointly optimizes its reconstruction loss with a self-supervised clustering loss. By minimizing a clustering loss iteratively, the batch effect in the embedding is reduced and cell type signal is improved^3^. The denoised gene expression values, computed from this embedding using a decoder, are then corrected for batch effects as well. To address the difficulty of batch correcting genes that are not highly variable, which suffer from a lower cell type signal-to-noise ratio, we designed CarDEC using a branching architecture that treats highly variable genes (HVGs) and the remaining genes, which we designate as lowly variable genes (LVGs), as distinct feature blocks.

CarDEC is unique among batch effect correction methods in that it implicitly corrects for batch effect through joint optimization of its dual objective function, rather than explicitly modeling batch effect using batch indicators as in methods such as MNN^5^ and scVI^4^. Moreover, it corrects batch effect both in the low-dimensional embedding space and the original gene expression space. CarDEC’s architecture is uniquely founded on the idea of treating HVGs and LVGs as different “feature blocks,” which enables CarDEC to use the HVGs to drive the clustering loss, while still allowing the LVG reconstructions to depend on the rich, batch corrected embedding learned from the HVGs, to help remove batch effect in the LVGs. Through comprehensive analyses on numerous datasets spanning different species and tissues with various degrees of complexities, we show that CarDEC is effective in removing complex batch effect and consistently outperforms scVI^4^, DCA^12^, MNN^5^, and scDeepCluster^13^ for both batch effect correction and clustering accuracy. We also show that with appropriate denoising and batch effect correction, the LVGs offer as much signal for clustering as the HVGs. Furthermore, the effective batch effect correction in gene expression offered by CarDEC allows it to reveal biologically anticipated marker genes in trajectory analysis that are otherwise obscured in the presence of batch effect by other methods.

## Results

### Overview of CarDEC and evaluation

An outline of the CarDEC workflow is shown in **Figure 1** and **Supplementary Figure 1**. CarDEC starts by data preprocessing and pretraining of an autoencoder using HVGs with a mean squared error reconstruction loss function. After pretraining, the weights learned from the pretrained autoencoder are transferred over to the main CarDEC model, which treats HVGs and LVGs as different feature blocks. The main CarDEC loss function is a weighted combination of the reconstruction losses for the HVGs and the LVGs, and a self-supervised clustering loss function driven by the HVGs. This combined loss function allows CarDEC to preserve local structure of the data during clustering^14^. By minimizing this self-supervised combined loss function, CarDEC not only improves the low-dimensional embedding for clustering, but the reconstructed genewise features, which are computed as a function of the low-dimensional embedding, is also denoised and batch effect corrected, leading to dramatically improved gene expression quality.

**Figure 1.**
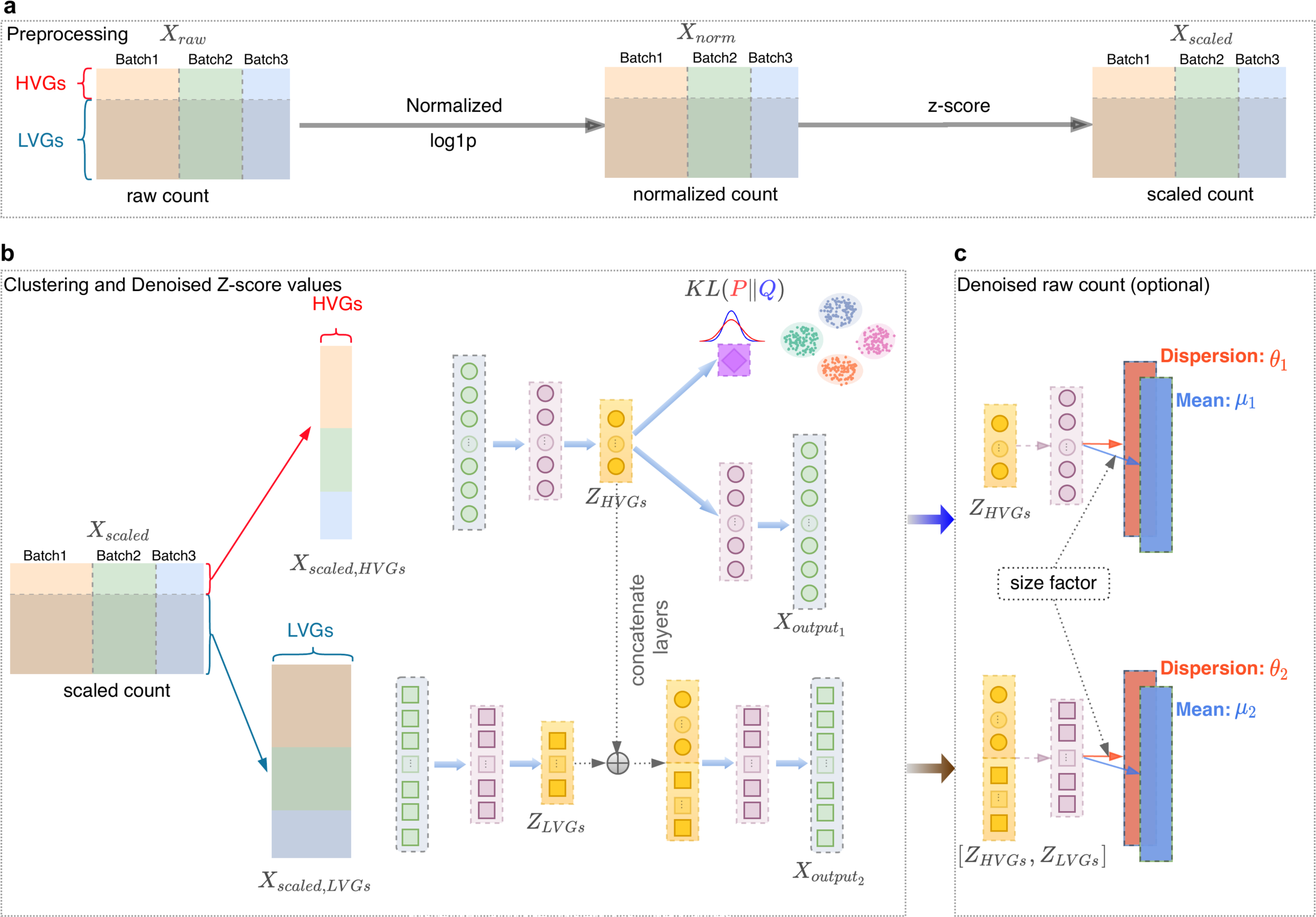
The workflow of CarDEC. The CarDEC workflow can be summarized in four steps that are depicted here: preprocessing, pretraining, denoising, and optionally, denoising on the count scale.

We evaluated CarDEC on a diverse set of challenging real datasets that range from human to mouse and have different flavors of batch effect. In our evaluations, we wish to assess two properties of CarDEC: 1) its ability to recover biological signals in the data, and 2) its ability to remove spurious technical signals driven by batch effect. An ideal method should strive to remove batch effect while maintaining true biological variations. We compared CarDEC with several state-of-the-art scRNA-seq methods for denoising, batch effect correction, and clustering. scVI^4^ and DCA^12^ are multi-use methods that provide denoised counts in the gene expression feature space, and also a low-dimensional embedding that can be used for tasks like clustering and visualization. scVI also attempts to correct for batch effect by conditioning on batch annotation when modeling the denoised counts from a zero-inflated negative binomial distribution. MNN^5^ is a batch correction method that merges batches in a pairwise manner and generates batch corrected gene expression on a cosine scale. scDeepCluster^13^ is a clustering method that also draws inspiration from the self-supervised clustering loss^14^.

To measure the degree of batch mixing, we examined the batchwise centroids before and after denoising and/or batch correction for each method by calculating a coefficient of variation (CV) metric. For each gene, the CV is calculated using the centroid of each batch. A higher value of CV corresponds to greater variation of gene expression among batches and less batch mixing, whereas a good batch effect removal method should drive the CV value close to zero.

### Application to human pancreatic islet data from four protocols

A unique feature of CarDEC is the branching architecture for both the HVGs and the LVGs. To demonstrate that this architecture is key in removing batch effect, we combined four datasets on human pancreas generated using Fluidigm C1^15^, SMART-seq2^16^, CEL-seq^17^, and CEL-seq2^18^. The branching architecture was designed with two objectives in mind. First, we wish to show that when correcting batch effect and denoising both the HVGs and the LVGs, using a branching model that treats these feature blocks differently improves the quality of denoised expression values relative to a naïve architecture that treats these feature blocks the same. Second, we hope to design a model architecture such that including the LVGs in the model does not worsen denoising and batch effect correction quality of the HVGs, relative to a naïve model that only denoises the HVGs and does not attempt to denoise LVGs.

As shown in **Figure 2**, the branching architecture posts significant performance boosts over the naïve architecture that treated all genes as the same feature block in the input. The branching architecture performed better for denoising both the HVGs (Adjusted Rand Index (ARI) of 0.93 over 0.72) and the LVGs (ARI of 0.83 over 0.67) relative to the naïve architecture (**Figure 2a,b**), underscoring the necessity of using the branching architecture to denoise all genes as efficiently as possible. We also observed that for the purpose of denoising and batch correcting only the HVGs, the branching architecture performed just as well as a naïve model that only included the HVGs and completely discarded the LVGs (ARI of 0.93 vs 0.94) (**Figure 2a,c**). This verifies that the branching architecture does not trade off denoising and batch correction effectiveness on the HVGs at all to denoise the LVGs. Additionally, the denoised counts from CarDEC showed considerably less batch effect compared to denoised expression from scVI and batch corrected expression from MNN (**Supplementary Figure 2**). We also noticed that the clustering accuracies are similar for denoised values and embedding for CarDEC, but the clustering accuracy is much lower when using denoised values as input than using embedding for scVI (**Supplementary Figure 3**). Strikingly, the ARI for scVI with denoised gene expression as input is even lower than that when using raw read counts as input (**Supplementary Figure 4**). This result suggests that batch effect correction in the gene expression space is much harder than in the embedding space, consistent with the findings of Lucken *et al*.^11^.

**Figure 2.**
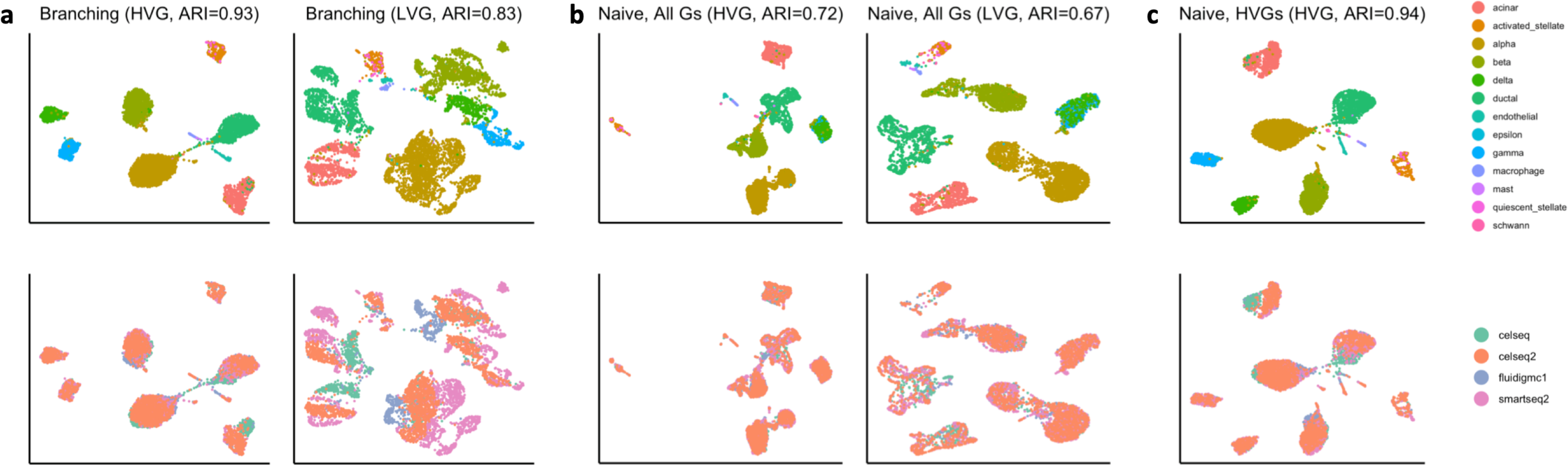
Justification for the branching architecture in CarDEC. The CarDEC API splits the input matrix into HVGs and LVGs and treats them separately with a “Branching” architecture as in Fig. 1. Alternatively, we can use a “Naïve” model for finetuning that treats all features the same regardless of gene expression variance, which consists of an autoencoder with a clustering loss in addition to the reconstruction loss. Here we demonstrate the utility of the Branching architecture. The HVGs and LVGs are clustered separately and the ARI of assignments is provided along with a UMAP plot. First row colored by cell type, second by scRNA-seq protocol. **a**, Clustering using denoised counts from the CarDEC Branching architecture. **b**, Clustering using denoised counts from the CarDEC Naïve architecture. All genes (HVGs and LVGs) were treated the same and denoised together. **c**, Clustering using denoised counts from the CarDEC Naïve architecture, but using HVGs only in the Naïve model. Since the LVGs were not included in this scenario, evaluation was only done for denoised expression for the HVGs.

### Application to macaque retina data with multi-level batch effect

After finalizing the CarDEC architecture, we next evaluated the performance of CarDEC on a macaque retina dataset^19^. This dataset poses a great challenge for batch effect correction and denoising because it features a strong, multi-level batch effect, with cells sequenced from two different regions, four different macaques, and thirty different samples (**Supplementary Figure 5**).

For the task of denoising and batch effect correction in the gene expression space, CarDEC was again the best performing method (**Figure 3, Supplementary Figures 6 and 7**), and the gap in performance between CarDEC and other methods is even larger on this more challenging dataset. CarDEC not only removed the multi-level batch effect but also preserved inter-cell type variation. Notably, the ARI for clustering using the LVG denoised and batch effect corrected counts from CarDEC is 0.98 (**Figure 3b**), which is as high as that using the HVGs (**Figure 3a**). As a comparison, the ARI is only 0.15 using the LVG raw counts as input for clustering (**Supplementary Figure 5**). This suggests that the denoising and batch correction in CarDEC substantially boosted the signal-to-noise ratio in the LVGs. Moreover, CarDEC’s genewise CVs are consistently the closest to zero, providing evidence that cells were mixed well by batch (**Figure 3c, Supplementary Figure 8**).

**Figure 3.**
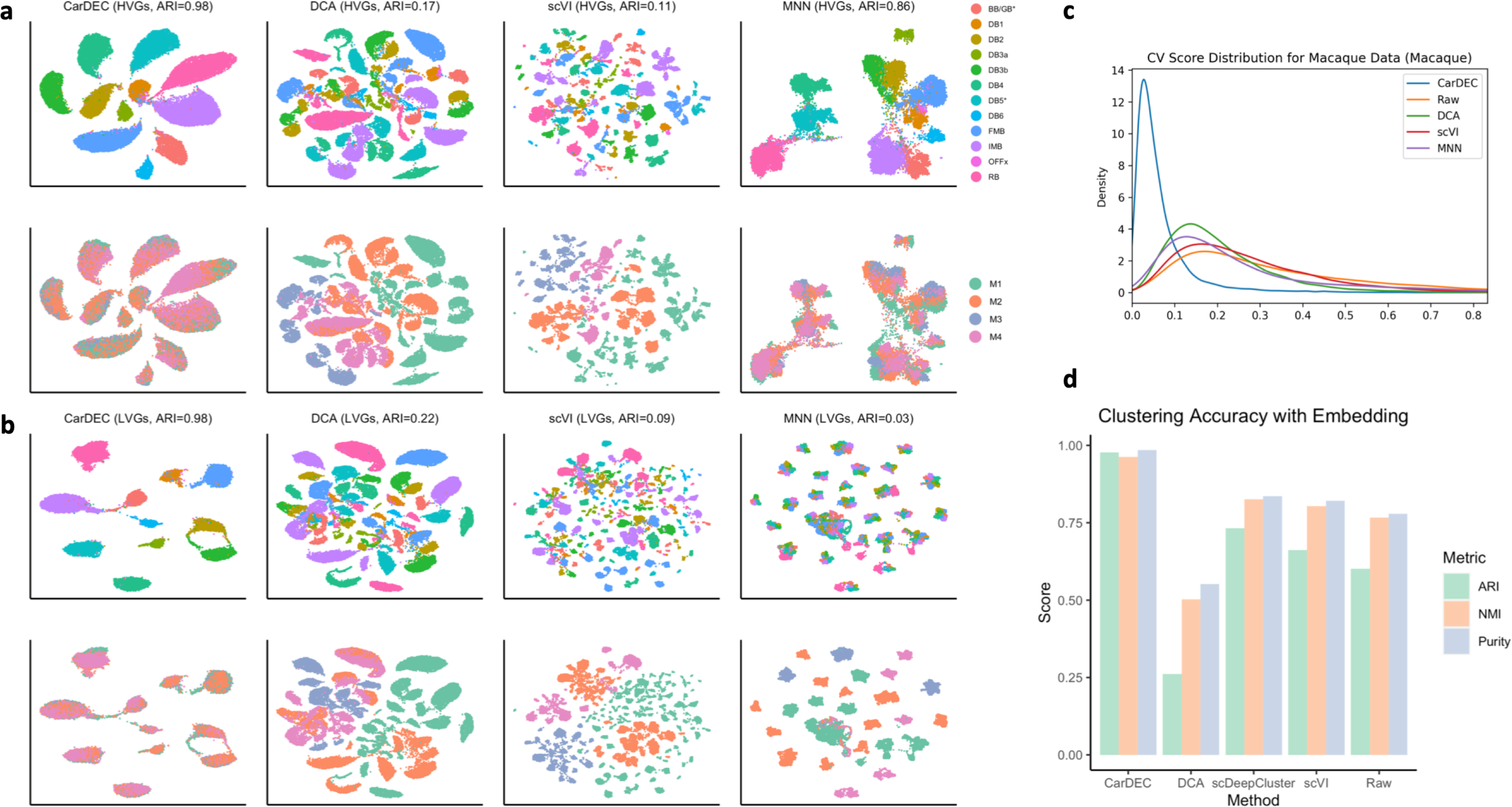
Comparison of different methods on the macaque retina dataset. **a**, UMAP embedding computed from the denoised HVG counts for each method. Top row colored by cell type; bottom colored by Macaque ID. UMAPs colored by region id and sample id provided in the supplement. Cells were also clustered with Louvain’s algorithm. **b**, UMAP embedding computed from the denoised LVG counts for each method. Figure legends are the same as those in **a. c**, Density plot of genewise coefficient of variation (CV) among batch centroids. Centroids computed with sample id as batch definition. CV plots with region id and macaque id as batch definition are provided in **Supplementary Figure 9. d**, Clustering accuracy metrics obtained using the embedding based methods to cluster the data, rather than running Louvain on the full gene expression space. Results for “Raw” were run using Louvain’s algorithm on the original HVG counts, to provide a baseline with which to compare embedding based clustering results to.

The other methods all struggled with batch effect in the denoised counts. scVI largely failed to correct batch effect: when using scVI batch corrected counts, the cells were separated primarily by batch rather than by cell type. For the LVGs, its ARI is even lower than that using the LVG raw counts as input for clustering (scVI 0.09 vs raw 0.15) (**Figure 3b, Supplementary Figure 5**). DCA had slightly higher ARIs than scVI for both the HVGs and the LVGs, although both are significantly lower than CarDEC (**Figure 3a,b**). MNN performed much better than scVI and DCA for batch correcting the HVG counts, achieving an ARI of 0.86 (**Figure 3a**). However, it still fell substantially short of CarDEC for this evaluation (CarDEC ARI 0.98). Looking more closely at the HVG UMAP plots, the batches were mixed less thoroughly with MNN than they were for CarDEC, and the cells were separated less by cell type indicating that MNN failed to completely recover cell type variation. This is further confirmed by the genewise CV density plot in which the MNN density curve is further away from zero than CarDEC (**Figure 3c**). For removing batch effects in the LVG counts, MNN again was the worst performing method because it removed nearly all biological variations, leaving only batch effects.

For the simpler task of clustering using embedding, existing methods did considerably better than they did at batch effect correction in the gene expression space but still fell short of CarDEC (**Figure 3d, Supplementary Figure 9**). CarDEC achieved an ARI nearly 1 for clustering using the embedding. scDeepCluster and scVI performed slightly better than Louvain’s algorithm using raw HVGs, but still fell short of achieving 0.75 ARI. DCA struggled on this dataset with an ARI of only 0.25.

### Application to mouse cortex and PBMC data from four protocols

We next compared different methods using a mouse cortex dataset^20^. This dataset poses the greatest challenge for batch correction and denoising on two fronts (**Supplementary Figure 10**). First, it exhibits very serious batch effects owing to the fact that cells were generated using four different scRNA-seq protocols. Furthermore, this dataset is heavily dominated by excitatory and inhibitory neurons, and the other cell types are rare, so preserving biological variation is especially imperative for detecting and analyzing these rarer subpopulations.

For the task of denoising and batch correcting the gene expression space CarDEC performed considerably better than the other methods (**Figure 4**). CarDEC performed the best at balancing between removing batch effect while preserving as much cell type variability as possible. The ARIs are similar when using the HVG denoised counts and the LVG denoised counts as input for clustering (**Figure 4a,b**). As a comparison, the ARIs are only 0.26 and 0.25 when using the HVG and the LVG raw counts as input for clustering, respectively (**Supplementary Figure 10**). The relatively low ARI when using the HVG raw count as input for clustering demonstrates the strong batch effect in this dataset. However, for this challenging dataset, using CarDEC denoised and batch corrected LVG counts, the ARI increased to 0.74, suggesting that CarDEC substantially boosted the signal-to-noise ratio in the LVGs by simultaneous denoising and batch effect removal. The genewise CVs for CarDEC are also the closet to zero among all methods (**Figure 4c**).

**Figure 4.**
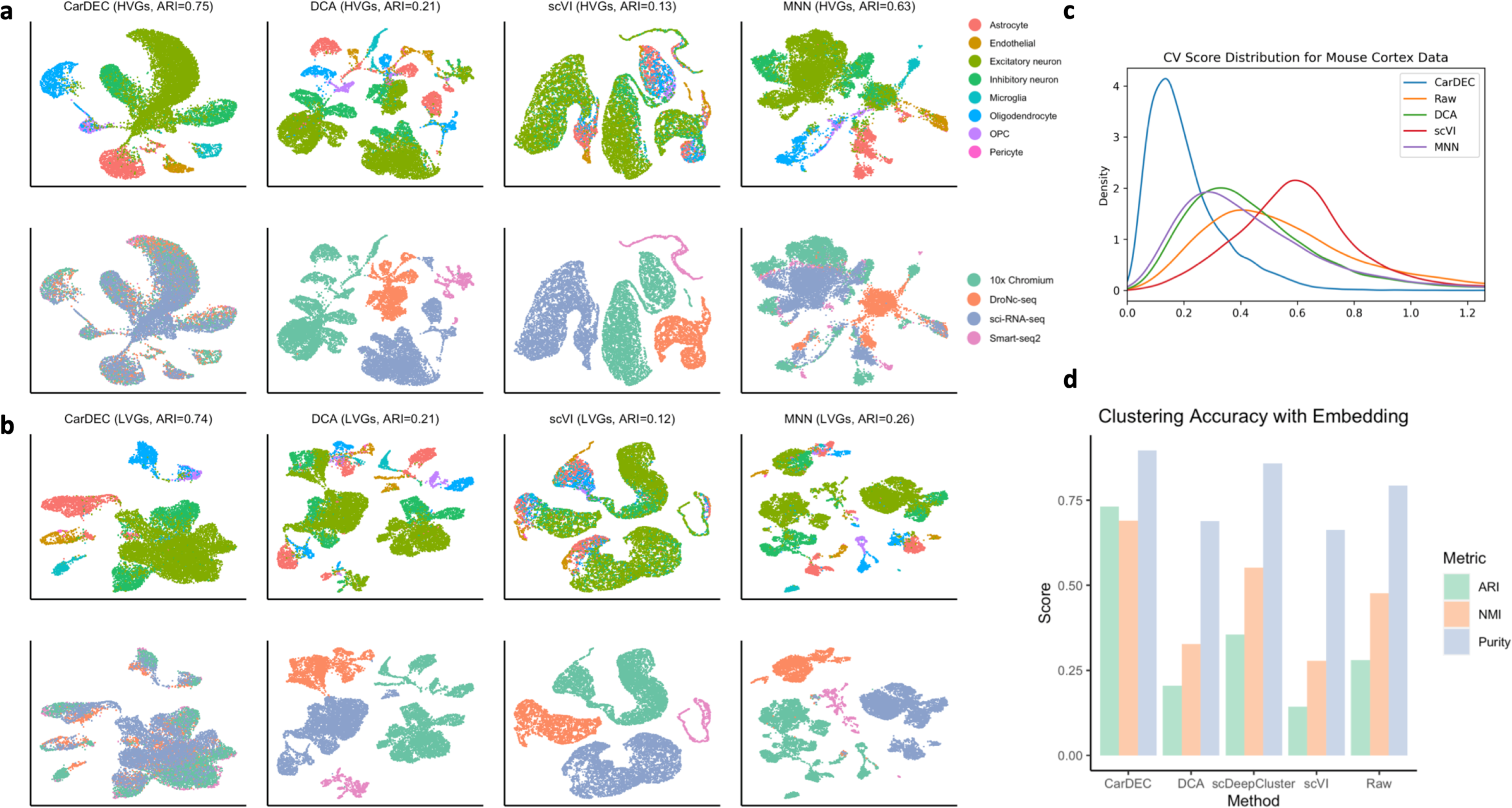
Comparison of different methods on the mouse cortex dataset. **a**, UMAP embedding computed from the denoised HVG counts for each method. Top row colored by cell type; bottom colored by batch. Cells are also clustered with Louvain’s algorithm, and resultant ARI is provided. **b**, UMAP embedding computed from the denoised LVG counts for each method. Figure legends are the same as those in **a. c**, Density plot of genewise coefficient of variation (CV) among batch centroids. **d**, Clustering accuracy metrics obtained using the embedding based methods to cluster the data, rather than running Louvain on the full gene expression space. Results for “Raw” were run using Louvain’s algorithm on the original HVG counts, to provide a baseline with which to compare embedding based clustering results to.

By contrast, DCA and scVI largely failed for this dataset (**Figure 4a,b**). For both the HVGs and the LVGs, DCA and scVI separated the cells purely by scRNA-seq protocol with no mixing of cells from different batches. After denoising using DCA and scVI, cell variation was driven entirely by batch, rendering the denoised counts ineffective for downstream analyses. Consistent with our previous evaluations, for batch correcting the HVGs, MNN was the second-best performer (**Figure 4a**), ahead of other methods by substantial margins but still lagging relative to CarDEC. MNN did not merge batches to the extent that CarDEC did and failed to preserve as much cell type variability, causing cell types to mix more. For removing batch effects in the LVGs, MNN did considerably worse than CarDEC and only slightly better than DCA and scVI (**Figure 4b**). It suffered from the same problems as DCA and scVI for the LVGs in that cell type variation was lost and all variability was driven by batch.

Even the simpler task of clustering the data using embedding was very difficult on this dataset (**Figure 4d, Supplementary Figure 11**). Both DCA and scVI performed poorly at this task, scoring lower ARIs than a straightforward application of Louvain’s algorithm to the raw data. scDeepCluster showed slightly better performance, but its ARI still fell below 0.4. For this task, CarDEC also was the clear leader, achieving an ARI of 0.73.

We also analyzed a dataset of human PBMCs from the same paper^20^ as the mouse cortex data. This dataset was similar to the cortex dataset: featuring eight batches spanning five scRNA-seq protocols and the results were largely the same: CarDEC was the best for denoising/batch correcting the HVGs and far away the best for denoising/batch correcting the LVGs (**Supplementary Figures 12-14**).

### Application to human monocyte data with pseudotemporal structure

We next show the utility of CarDEC for improving trajectory analysis for cells with pseudotemporal structure. We analyzed a scRNA-seq dataset generated from monocytes derived from human peripheral blood mononuclear cells by Ficoll separation followed by CD14- and CD16-positive cell selection^3^. This dataset includes 10,878 monocytes from one healthy subject. The cells were processed in three batches from blood drawn on three different days. Although monocytes can be classified as classical (CD14^++^/CD16), intermediate (CD14^++^/CD16^+^), and nonclassical patrolling (CD14^-^/CD16^++^) subpopulations based on surface markers, our previous analysis based on scRNA-seq data indicates that these cells show continuous transcriptional characteristics and trajectory analysis is an appropriate approach to characterize them^3^. This dataset has strong batch effect (**Supplementary Figure 15**). To reconstruct the trajectories of these cells, for each method, we first denoised and/or batch corrected the gene expression matrix, which was then fed into Monocle 3^21^ to estimate the pseudotime of each cell.

**Figure 5a** shows that CarDEC yields far and away the best pseudotime analysis results with cells from the three batches well mixed, and a clear pseudotemporal path emerged. The batchwise density plots show that the three batches have similar pseudotime distributions, suggesting that CarDEC successfully removed batch effect. The plots of *FCGR3A* (known marker gene for nonclassical monocytes) and *S100A8* (known marker gene for classical monocytes) gene expression also showed expected patterns (**Supplementary Figure 16**). There are two key points of evidence from these marker gene plots suggesting that CarDEC recovered biological signal. First, for each marker gene, the expression levels are virtually identical across batches for all pseudotime points, which indicates that batch effect was removed for each gene expression and pseudotime relationship. Also, *FCGR3A* gene expression decreases monotonically with pseudotime, while *S100A8* expression increases monotonically with pseudotime. This is exactly the kind of behavior we expect from these marker genes. Since *FCGR3A* and *S100A8* are markers for the nonclassical and classical monocytes, respectively, we expect a good pseudotime analysis to segment the monocytes from nonclassical to classical (or vice versa) and for *FCGR3A* and *S100A8* expressions to be monotonic function of pseudotime with opposite trends. By denoising and batch correcting gene counts, CarDEC successfully mixed batches and recovered biological signal down to the individual marker gene level.

**Figure 5.**
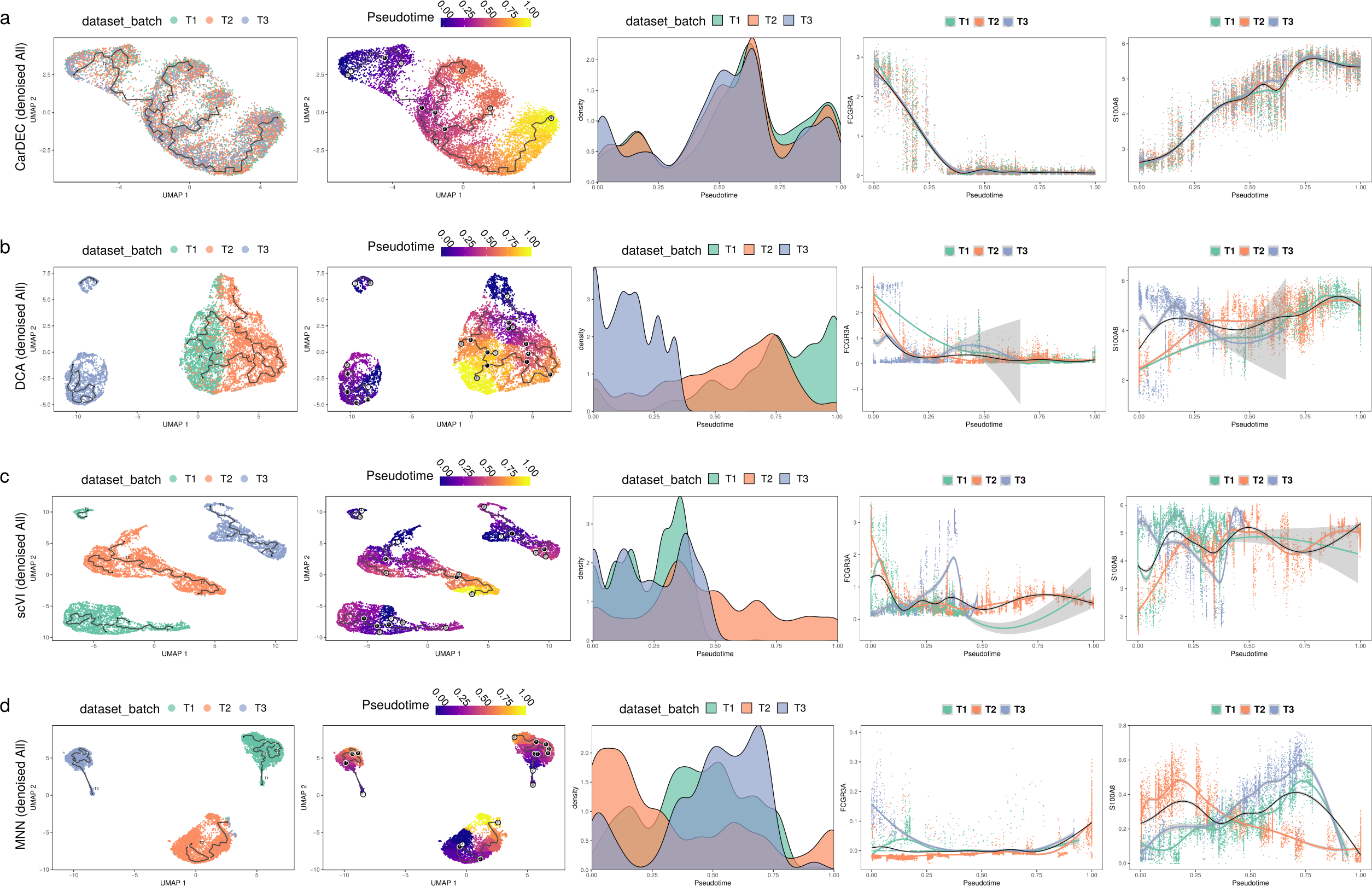
Comparison of different methods for pseudotime analysis in the human monocyte data. The analysis is for Monocytes derived from three technical replicates from the same subject. For each method, the full dataset was denoised/batch corrected and then fed to Monocle 3 for pseudotime analysis. We show the UMAP embedding colored by batch (column 1) and estimated pseudotime (column 2). We also visualize the kernel density distribution of pseudotime by batch (column 3) and plot the distributions of marker genes *FCGR3A* and *S100A8* against pseudotime (columns 4 and 5, respectively). **a**, Pseudotime analysis when using denoised/batch corrected gene expression matrix from CarDEC as input. **b**, Pseudotime analysis when using denoised gene expression matrix from DCA as input. **c**, Pseudotime analysis when using denoised/batch corrected gene expression matrix from scVI as input. **d**, Pseudotime analysis when using batch corrected gene expression matrix from MNN as input.

By contrast, no other methods were able to achieve CarDEC’s success in improving pseudotime analysis. DCA (**Figure 5b**), scVI (**Figure 5c**), and MNN (**Figure 5d**) all failed to mix batches in the UMAP embedding from Monocle 3, and the pseudotime distribution varied across batches for all three methods. Furthermore, neither of the marker genes show strong monotonic trends as a function of the pseudotime, and for each marker gene, the relationship between expression and pseudotime varied by batch. These issues suggest that DCA, scVI, and MNN’s failures to correct for batch effects confounded biological signal and obscured signals from canonical markers of established subpopulations, *FCGR3A* and *S100A8*, as marker genes using these approaches.

There are other approaches to using these denoising and batch correction methods for pseudotime analysis. For example, one can subset the denoised and batch corrected matrix to include only the HVGs and then feed this into Monocle 3 (**Supplementary Figures 17 and 19**). Alternatively, one can use the embedding from CarDEC, scVI, or DCA as the reduced dimension space to build the Monocle 3 pseudotime graph (**Supplementary Figures 18 and 20**). In both of these other cases, the conclusions are largely the same, CarDEC is far and away the best method for improving pseudotime analysis.

Next, we examined whether the denoised and batch corrected gene expression values can help improve gene expression quality for biological discovery. We focused our analyses on 61 transcription factors (TFs) that were expressed in the monocyte data and also found to be differentially expressed among classical, intermediate, and nonclassifcal monocytes by Wong *et al*.^22^. Among these 61 TFs, 23 were selected as HVGs and the remaining 38 were designated LVGs. **Figure 6a** shows that the CarDEC denoised gene expression revealed a gradual decreasing trend from nonclassical to classical for TFs that are known to be highly expressed in nonclassical monocytes, e.g., *TCF3L2, POU2F2, CEBPA*, and *HSBP1*. We also observed expected gene expression increase from nonclassical to classical for TFs that are known to be highly expressed in classical monocytes, e.g., *NEF2, CEBPD, GAS7*, and *MBD2*. Notably, some of the TFs with these expected expression patterns were not selected as HVGs, suggesting that denoising and batch correction in CarDEC helped recover the true biological variations. By contract, when using raw UMI counts as input, the heatmap did not reveal any meaningful biological patterns even for those TFs that were selected as HVGs (**Figure 6b**).

**Figure 6.**
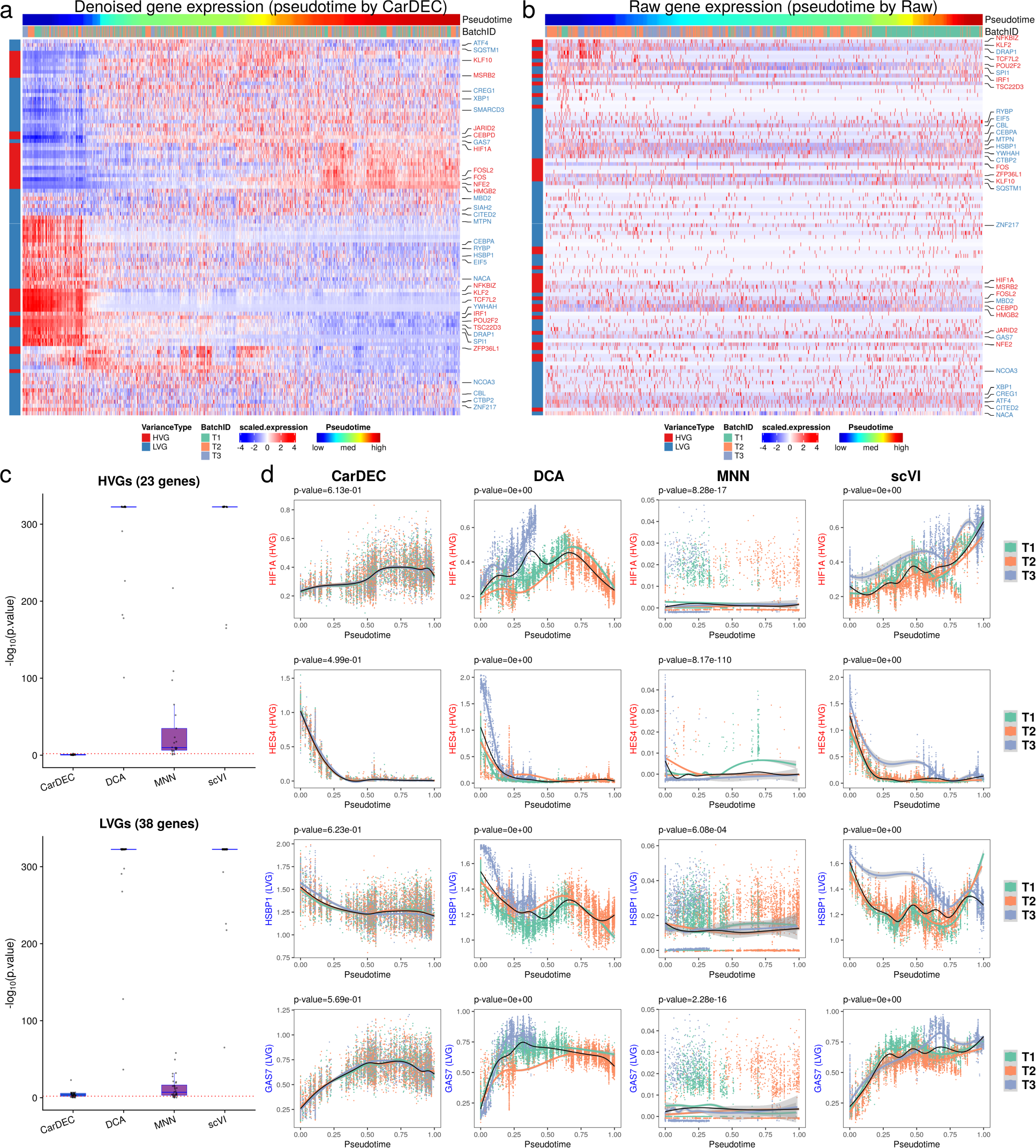
Comparison of different methods for differential expression analysis of transcription factors in the human monocyte data. **a**, Heatmap of scaled gene expression for CarDEC. Pseudotime was inferred based on embedding obtained from CarDEC using Monocle 3. **b**, Heatmap of scaled raw UMI counts. Pseudotime was inferred based on embedding obtained from the scaled raw UMI counts using Monocle 3. **c**, p-values obtained from differential expression analysis among the three batches over pseudotime. For each method, the pseudotime was inferred based on embedding obtained from the corresponding method. The red dotted line corresponds to p-value = 0.01. The top panel is for the 23 HVG TFs and the bottom panel is for the 38 LVG TFs. **d**, Denoised and batch corrected gene expression for CarDEC, denoised gene expression for DCA, batch corrected gene expression for MNN, and denoised and batch corrected gene expression for scVI over pseudotime for four selected TFs, *HIF1A, HES4, HSBP1*, and *GAS7*. For each method, the pseudotime was inferred based on embedding from the corresponding method.

An important task in trajectory analysis is to identify genes whose expression values change over pseudotime and whether the expression patterns are different between conditions (e.g., healthy vs diseased) over pseudotime. Avoiding generating false positive results is critical as failure of doing so may lead to follow-up of a wrong signal. Since the three batches were obtained from the same subject, we do not expect to detect significant gene expression differences over pseudotime among them. To this end, we performed differential expression analysis and compared the distribution of gene expression changes over pseudotime across the three batches. We performed hypothesis tests using the ‘*gam*’ function in R package *mgcv* and tested whether gene expression patterns for the three batches are significantly different over pseudotime. **Figure 6c** shows the p-values from this differential expression analysis for each method. CarDEC is clearly much more effective in removing batch effect than DCA, MNN, and scVI. The median -log10 p-value for CarDEC is 3.70, whereas the median -log10 p-values for DCA, MNN, and scVI are 322, 7.08, and 323, respectively. These results clearly indicate that failure to correct for batch effect could lead to a severe inflation of false positive results. **Figure 6d** shows four selected TFs, where the denoised and batch corrected gene expression for CarDEC agreed well among the three batches, further confirming the effectiveness of CarDEC in removing batch effect in the gene expression space. Gene expression plots for the remaining 57 TFs are shown in **Supplementary Figure 21**.

### CarDEC is scalable to large dataset

As the scale of scRNA-seq continues to grow, it becomes increasingly important for a method to be scalable to large datasets. To evaluate the scalability of CarDEC, we leveraged a dataset of 104,694 human fetal liver cells^23^. Since we are principally interested in the problem of denoising and batch correcting in the full gene expression space, we retained all 21,521 genes after initial filtering for this analysis. For CarDEC we benchmarked two variations: a version that provides only denoised/batch corrected expression in the Z-score space (CarDEC Z-score) and a version that provides denoised/batch corrected expression in the count space (CarDEC Count).

We evaluated the runtime needed to process 10%, 20%, 40%, 60%, 80%, and 100% of cells in the human fetal liver dataset for CarDEC, scVI, DCA, and MNN. All evaluations were done on a 2019 edition MacBook Pro with 2.4 GHz 8-Core Intel Core i9 CPU and 32 GB of memory. CarDEC, DCA, and scVI were all trained with early stopping to halt training upon convergence. The results are shown in **Supplementary Figure 22**. Both versions of CarDEC as well as DCA scaled approximately linearly with the number of cells and all three of these methods finished the analysis in less than 3.5 hours. scVI compared less favorably, as it took almost 11 hours to process the full dataset. MNN, on the other hand, has serious scalability issues, which is consistent with a recent benchmarking study^5^. It took over 12 hours to analyze 20% of the dataset, and over 47 hours to analyze 40% of the dataset. We could not run MNN in under 48 hours using more than 40% of the data.

## Discussion

We developed CarDEC, a joint deep learning model, that removes batch effects not only in the low-dimensional embedding space, but also across the entire gene expression space. As demonstrated in our evaluations and a recent benchmarking study^11^, it is considerably harder to correct for batch effect in the gene expression space than in the embedding space, and especially hard to correct for batch effect in LVGs, which constitute the majority of the transcriptome. CarDEC was built to tackle these challenges. To remove batch effect in the gene expression space, we minimize a loss function that combines clustering and reconstruction losses. The self-supervised clustering loss, driven by HVGs, regularizes the embedding and removes batch effects in the embedding. The rich, batch corrected embedding is then used to compute an effectively batch corrected representation in the original gene expression space. To address the difficulty associated with batch correcting LVGs, we implemented a branching architecture, where embeddings are computed separately for HVGs and LVGs and where only the HVG embedding is used to compute the clustering loss. Using the pancreatic islet datasets generated from four scRNA-seq protocols, we demonstrated that this branching architecture substantially improved batch effect removal on both the HVG and LVG gene expression spaces, as compared to the naïve architecture.

Across a variety of datasets, with batch effects spanning multiple complexities in level and strength we demonstrated that CarDEC consistently led in its ability to remove batch effects. CarDEC was consistently the best for removing batch effects in all capacities: in the embedding space, the HVG expression space, and the LVG expression space. In particular, CarDEC is the only method capable of batch correcting the LVGs. We showed that with appropriate denoising and batch correction, the LVGs offer as much signal for clustering as the HVGs, suggesting that CarDEC has substantially boosted the amount of information content in scRNA-seq. We also demonstrated that by batch correcting gene expression counts, CarDEC improved pseudotemporal analysis of human monocytes, an example of how batch correction can be used to improve downstream analyses.

Current scRNA-seq studies often include a large number of cells generated from many samples, across multiple conditions, and possibly using different protocols. Removing batch effect is critical for data integration. Since CarDEC provides efficient batch correction in the full gene expression space, it can be used to for a wide array of analyses to facilitate biological discovery. Harmonized counts in the gene expression space can be used to estimate unbiased, batch corrected log fold changes, which can be used to identify marker genes for different cell types. These counts can also be used to reconstruct trajectories and identify genes showing pseudotemporal patterns. Lastly, CarDEC is computationally fast and memory efficient, making it a desirable tool for analyses of complex data in large-scale single-cell transcriptomics studies.

## Acknowledgements

This work was supported by the following grants: R01GM125301 (to M.L.), R01EY030192 (to M.L.), R01EY031209 (to M.L.), R01HL113147 (to M.L. and M.P.R.), and R01HL150359 (to M.L. and M.P.R.). We thank Sean Simmons and Jiarui Ding for their help on the mouse cortex and human PBMC data analysis.

## Author contributions

This study was conceived of and led by M.L.. J.L. designed the model and algorithm, implemented the CarDEC software, and led data analysis with input from M.L. and X.L.. X.L. led data analysis for the human monocyte data and designed the workflow figure. D.W., G.H., and K.W. participated the early stage of algorithm design and testing. Y. Z. provided input on memory management and analysis for the human monocyte data. L. U. provided input on the model and algorithm design. H.Z. and M.P.R. provided input on the human monocyte data analysis. J.L. and M.L. wrote the paper with feedback from all coauthors.

## Methods

The CarDEC workflow (**Figure 1, Supplementary Figure 1**) involves four steps: preprocessing, pretraining, gene expression denoising in Z-score space, and (optionally) denoising in count space. Below we briefly describe each of these steps. Details of the implementation is described in **Supplementary Note 1**, and the hyperparameters of CarDEC are shown in **Supplementary Table 1**.

### Step 1: preprocessing

We first remove any cells expressing less than 200 genes, and then remove any genes expressed in less than 30 of the remaining cells. Let **X** be an *n* × *p* gene count matrix with *n* cells and *p* genes after filtering. The gene expression values are normalized. In the first step, cell level normalization is performed in which gene expression for a given gene in each cell is divided by the total gene expression across all genes in the cell, multiplied by 10,000, and then transformed to a natural log scale. In the second step, gene level normalization is performed in which the cell level normalized values for each gene are standardized by subtracting the mean and dividing by the standard deviation across all cells within the same batch for the given gene. Highly variable genes (HVGs) are selected based on the log-normalized counts using the approach introduced by Stuart and Butler^24^ and implemented in the “pp.highly_variable_genes” function with “batch_key” parameter in the Scanpy package (version >=1.4)^25^. The remaining genes that are not selected as HVGs are considered lowly variable genes (LVGs). We note that many of the LVGs still show cell-to-cell variability, and are useful for clustering analysis after appropriate denoising and batch effect correction. We select 2,000 HVGs for all analyses in this paper.

### Step 2: pretraining using the HVGs

The pretraining step is a straightforward implementation of an autoencoder. Let *p*_*HVG*_ be the number of HVGs selected in Step 1, and **Y**_*HVG*_ be the corresponding *n* × *p*_*HVG*_ matrix of normalized expression, subsetted to include only the HVGs. Define a standard autoencoder for Y_*HVG*_ with encoder and decoder represented by *f*_*E,HVG*_;(*W* _*E,HVG*_) and *f* _*D,HVG*_(; *W* _*D,HVG*_), respectively. The weights *W* _*E,HVG*_ and *W* _*E,HVG*_ are randomly initialized using the glorot uniform approach, and are tuned during pretraining. We use the tanh activation for the output of the encoder, and the linear activation function for the output of the decoder. For all intermediate hidden layers in the encoder and decoder, we use the ReLu activation function. The autoencoder is pretrained with mean squared error loss using minibatch gradient descent with the Adam optimizer^26^.

### Step 3: denoising Z-scores

In this step, we use an expanded, branching architecture to accommodate LVGs, and introduce a clustering loss that regularizes the embedding and improves batch mixing and denoising especially in the gene space. Let *p*_*LVG*_ be the number of LVGs selected in Step 1, and **Y**_*LVG*_ be the corresponding *n* × *p* _*LVG*_ matrix of normalized expression, subsetted to include only the LVGs, and ***y*** _*i,HVG*_ and ***y*** _*i,LVG*_ be the vectors of HVGs and LVGs, respectively in cell *i*. We retain the encoder and decoder mappings for HVGs, *f* _*E,HVG*_(· ; *W* _*E,HVG*_)and *f* _*D,HVG*_ (· ; *W* _*D,HVG*_) from Step 2, including the learnt weights *W* _*E,HVG*_ and *W* _*D,HVG*_. We introduce a clustering layer that takes the HVG embedding ***z***_*i,HVG*_ = *f* _*E,HVG*_ (*y*_*i,HVG*_; *W* _*E,HVG*_) as input and returns for each cell a vector of cluster membership probabilities for *h* clusters, where *h* is a user specified number. For this clustering layer, we introduce an *h* × *d* matrix of trainable weights/cluster centroids **M**, where the *j*^*th*^ row of **M** is a cluster centroid ***μ***_*j*_, and *d* is dimension of the embedding.

To initialize **M**, we run Louvain’s algorithm on the embeddings { ***z***_*i,HVG*_: *i* ∈ {1, 2, …, *n*}} learned from the pretrained autoencoder, and find the cluster centroid for each cluster. The clustering layer computes a vector of cluster membership probabilities for cell *i*, denoted by *q*_*i*_. Let *q*_*ij*_, the *j*^*th*^ element of *q*_*i*_, denote the probability that cell *i* belongs to cluster *j*. Then the membership probabilities are computed using a *t*-distribution kernel as follows,

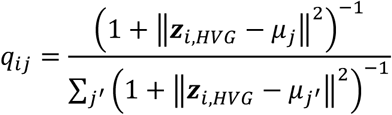

Since we do not have cell type labels in an unsupervised analysis, we create “pseudo-labels” that can be used in place of real labels for optimizing clustering weights. Inspired by Xie *et al*.^27^, these pseudo-labels are computed from the membership probabilities *q*_*ij*_ as follows,

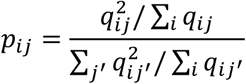

Let ***p***_*i*_ be an *h*-dimensional vector whose *j*^*th*^ element is *p*_*ij*_. Then the clustering loss for cell *i* is defined as the following Kullback–Leibler divergence (KLD),

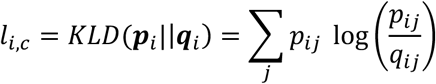

This loss is a component of the total loss defined later. Since it takes the embedding vectors ***z***_*i,HVG*_ as input, minimizing this objective function can refine the embedding and help to remove batch effects from denoised counts computed using this embedding as input.

We also introduce encoder and decoder mappings *f* _*E,LVG*_ (· ; *W* _*E,HVG*_) and *f* _*D,HVG*_ (· ; *W* _*D,HVG*_) to address the problem of denoising and batch correction for the LVGs. Unlike the HVG decoder *f* _*D,HVG*_, the LVG decoder *f* _*D,HVG*_ (·; *W* _*D,HVG*_) does not map the low-dimension embedding ***z***_*i,LVG*_ alone to reconstruct 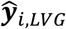 in the original *p*_*LVG*_ -dimension space. Rather, we concatenate the HVG and LVG embeddings together, and feed the combined vector [***z***_*i,HVG*_ ***z***_*i,LVG*_] into the decoder to denoise and batch correct LVG expression in the original *p* _*LVG*_ -dimension space. That is,

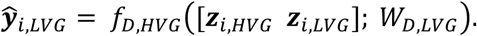

This concatenated embedding is critical because it allows CarDEC to only use the high signal-to-noise ratio HVGs to drive the clustering loss, while still using the rich, batch corrected embedding that is refined using this clustering loss to denoise and batch correct LVGs. The activation functions for the encoder and decoder of the LVGs are similarly defined as the autoencoder in Step 2.

To train this branching model, we first introduce two reconstruction losses, one for the HVGs and one for the LVGs computed as follows for cell *i*,

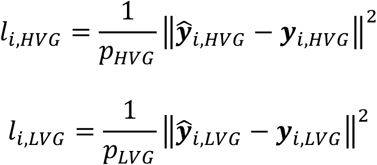

Then the total loss is calculated as a multi-component loss function as follows,

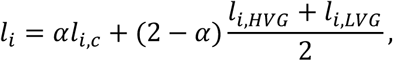

where *α* is a hyperparameter ranging from 0 to 2 that balances reconstruction loss with clustering loss. We set *α* at 1 as default value. The total loss is minimized in an iterative fashion until certain convergence criteria are satisfied.

### Step 4: denoising gene expression counts

In Step 3, the denoised expression values obtained from the decoder are on a Z-score scale and are not naturally comparable to raw UMI counts. To remedy this, we offer an optional downstream modeling step that provides denoised expression values on the original count scale. This strategy involves finding mean and dispersion parameters that maximize a negative binomial likelihood. We choose the negative binomial distribution because previous studies have shown that UMI counts are not zero-inflated, and negative binomial fits the data well^28-30^.

After the training in Step 3, we have obtained batch corrected low-dimension embeddings, ***z***_*i,HVG*_ and ***z***_*i,LVG*_ for each cell *i* from the fine-tuned HVG and LVG encoders. We will use two separate neural networks to maximize the negative binomial losses: one for the HVGs and one for the LVGs. These models are completely separate from one another but are trained almost identically with only minor differences. The goal is to map the embeddings into the full gene space to obtain mean and dispersion parameters for each gene. Without loss of generality, we use the HVGs as an example to illustrate how the neural network is built.

The vector of genewise means ***μ*** _*i,HVG*_ and vector genewise dispersions ***θ*** _*i,HVG*_ are given below,

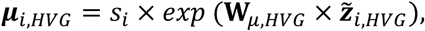

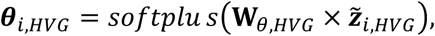

where *s*_*i*_ is the size factor for cell *i*, ***W*** _*μ,LVG*_ and ***W*** _*θ,HVG*_ are trainable weight matrices, and *exp* and *softplus* are activation functions that are applied elementwise. For each gene *j* in cell *i*, we compute the negative log likelihood of the negative binomial distribution as

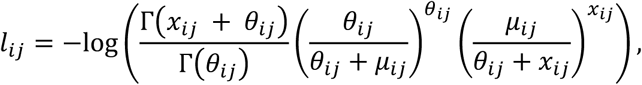

where *x*_*ij*_ is the original count in HVG gene *j* for cell *i, μ*_*ij*_ and *θ*_*ij*_ are the *j*^*th*^ elements of ***μ***_*i,HVG*_ and ***θ*** _*i,HVG*_, respectively. For the HVG count model, the full loss for cell *i* is then 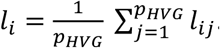. The loss for the LVG count model can be similarly defined. Both the HVG and LVG count models are trained using their own early stopping and learning rate decay convergence monitoring.

### Evaluation of batch effect removal in the gene expression space and the embedding space

Here, we briefly describe the workflow to evaluate batch effect removal and comparison between different methods (details see **Supplementary Notes 2 and 3**). First, we evaluated the performance of different methods in removing batch effect in the gene expression space. For this evaluation, we considered CarDEC, scVI, DCA, and MNN. We ran all denoising/batch correction methods on the full data matrix and then present clustering results for denoised HVGs and denoised LVGs separately by subsetting the HVGs and LVGs from the full denoised/batch corrected expression matrix. The subsetted matrix that includes only the HVGs (or LVGs) is then passed down to the Louvain’s clustering algorithm. All steps in this workflow are identical for both the HVGs and the LVGs, and all methods used the same HVGs and LVGs as input for clustering. Furthermore, on a given dataset, we benchmarked all methods with the same number of clusters. Second, we evaluated batch effect removal for the embedded representations of scRNA-seq. For this evaluation we considered CarDEC, scVI, DCA, and scDeepCluster. We excluded MNN since it has no embedding functionality. We also included “raw” as a control method for comparison, which is just subsetting the raw data to include only the HVGs, and then running the clustering workflow.

### Coefficient of variation (CV) analysis

To measure batch mixing, we examined the batchwise centroids before and after denoising. Let **X**′ be a matrix of gene expression counts (including both HVGs and LVGs). **X**′ can be the matrix of raw counts, or the denoised/batch corrected counts from any of CarDEC, scVI, DCA, or MNN. For CarDEC, we only considered denoised counts, not denoised expression in the Z-score space. If **X**′ consists of MNN corrected expression, then we did not preprocess the data since MNN denoised expression is on a cosine scale. In the case of MNN, a fraction of expression counts can be negative, which poses difficulties when computing coefficients of variation. To circumvent this issue, any MNN expression values that are negative were truncated to zero for the CV analysis. For all other methods we have denoised expression in the non-negative count space, so we performed cell normalization and log normalization on **X**′, exactly in the same way described in the Step 1 (preprocessing) of CarDEC.

Let *x*_*ij*_′ be the expression value in gene *j* of cell *i* in **X**′. Let *S*_5_ be a set of integers defined such that *i* ∈ *S*_*b*_ if and only if cell *i* was sequenced from batch *b*. Furthermore, let 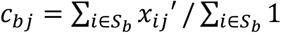 be the centroid (mean expression) of batch *b* for gene *j*. Let *C* _*j*_ = {*c*_*bj*_} be the set of batch centroids for gene *j*. If we have *B* batches, then this will be a set of *B* numbers. We define the CV for gene *j* to measure the degree of batch mixing as follows:

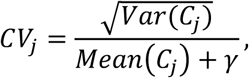

where *γ* is a small number included to guarantee computational stability for very lowly expressed genes. We set *γ* = 10^−12^. A higher value of *CV*_*j*_ corresponds to greater variation among batches and less batch mixing, and a good batch effect removal method should drive *CV*_*j*_ closer to zero. Normalizing by mean expression adjusts the CV for how highly expressed the gene is, so that CVs from more highly expressed genes are comparable to CVs from less highly expressed genes.

### Evaluation metrics for clustering

For all of our benchmark datasets, we used the cell type labels reported in the original papers as the gold standard. The clustering performance of each method was mainly evaluated using the adjusted rand index (ARI), calculated as below,

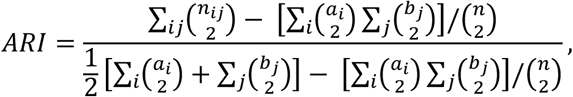

where *n*_*ij*_ is the number of cells in both cluster *i* from the cluster assignments obtained when benchmarking and in cell type *j* according to the gold standard cell type labels. *a*_*i*_ is the total number of cells in cluster *i* from the cluster assignments obtained when benchmarking, *b*_*j*_ is the total number of cells in cell type *j* according to the gold standard cell type labels from the original study, and *n* is the total number of cells. Additionally, we also computed normalized mutual information (NMI) and purity, calculated as below,

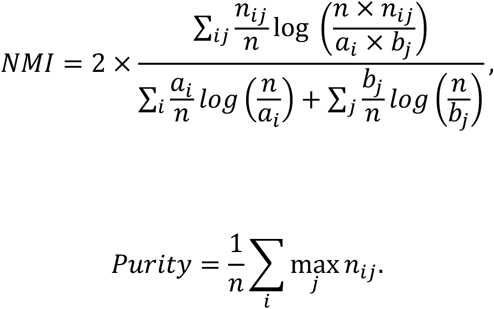

## Data availability

We analyzed multiple published scRNA-seq datasets, which are available through the accession numbers reported in the original papers. 1) Human pancreatic islet data: CelSeq (Gene Expression Omnibus GSE81076), CelSeq2 (Gene Expression Omnibus GSE85241), Fluidigm C1 (Gene Expression Omnibus GSE86469), and SMART-Seq2 (Array Express E-MTAB-5061); 2) Bipolar cells from mouse retina (Gene Expression Omnibus GSE81904); 3) Bipolar cells from macaque retina (Gene Expression Omnibus GSE118480); 4) mouse cortex data (Single Cell Portal SCP425); 5) human PBMC data (Single Cell Portal SCP424); 6) human monocyte data GEO (GSE146974); 7) human fetal liver data (Array Express E-MTAB-7407). Details of these datasets were described in **Supplementary Table 2**.

## Software availability

An open-source implementation of the CarDEC algorithm can be downloaded from https://github.com/jlakkis/CarDEC

## Life sciences reporting summary

Further information on experimental design is available in the Life Sciences Reporting Summary.

